# Proteolytic processing of the Marburg virus glycoprotein depends on Sec61β and is required for cell entry

**DOI:** 10.1101/2025.06.26.660697

**Authors:** Katharina E. Decker, Anke-Dorothee Werner, Markus Hoffmann, Heike Hofmann-Winkler, Qi-Yin Chen, Lu Zhang, Pamela Stomberg, Sabine Gärtner, Amy Kempf, Inga Nehlmeier, Hendrik Luesch, Torsten Steinmetzer, Eva Böttcher-Friebertshäuser, Nabil G. Seidah, Stefan Pöhlmann, Michael Winkler

## Abstract

Ebola and Marburg virus (EBOV, MARV) cause severe disease and therapeutic options are urgently needed. The Sec61 translocon facilitates ER import of viral glycoproteins (GPs) and may represent a therapeutic target. Here, we report that the Sec61 subunit Sec61β, although dispensable for GP expression, is required for proteolytic cleavage of MARV- but not EBOV-GP and that an intact furin motif is essential for robust cell entry of Marburg- but not Ebolaviruses. Further, MARV- but not EBOV-GP was cleaved by the furin-related enzyme SKI-1, for which a cleavage motif was identified in silico, and cleavage by SKI-1 was impaired in *SEC61B*-KO cells. In addition, Sec61β was required for normal N-glycosylation of MARV-GP and mutation of a sequon (N563D) abrogated cleavage. Finally, the absence of Sec61β modestly, and blockade of Sec61 via apratoxin S4 markedly, inhibited EBOV and MARV infection. These results reveal a differential protease dependence of MARV and EBOV and identify Sec61 as a potential therapeutic target.

**Author summary:** The filoviruses Ebola virus (EBOV) and Marburg virus (MARV) spread from animals to humans and can cause deadly outbreaks. These viruses rely on a surface glycoprotein (GP) for infection, which is processed by the enzyme furin in infected human cells. Cleavage of EBOV-GP was thought to be non-essential for infection. However, using lab models for filovirus entry into cells, we discovered that MARV, unlike EBOV, needs this cleavage step to infect cells efficiently. We also found that the host cell protein Sec61β is necessary for proteolytic processing and glycosylation of MARV-GP but not EBOV-GP. In addition, we showed that another cellular enzyme, SKI-1, can process MARV- but not EBOV-GP. Finally, we found that removing Sec61β or blocking Sec61 activity reduced infection by both viruses. These findings show key differences in how the two viruses interact with host cells and suggest that targeting Sec61 could be a promising new strategy to fight Ebola and Marburg virus infections.

## Introduction

Several African filoviruses within the genera *Orthoebolavirus*, including Ebola virus (EBOV, now termed *Orthoebolavirus zairense*), Sudan virus (SUDV, now termed *Orthoebolavirus sudanense*), Tai Forrest virus (TAFV, now termed *Orthoebolavirus taiense*), Bundibugyo virus (BDBV, now termed *Orthoebolavirus bundibugyoense*), and *Orthomarburgvirus*, including Marburg virus (MARV, now termed *Orthomarburgvirus marburgense*) as only species, are transmitted from bats to humans and can cause severe disease in afflicted patients [1–3]. In contrast, the Asian filovirus Reston virus (RESTV, now termed *Orthoebolavirus restonenese*) is believed to be apathogenic in immunocompetent persons [4, 5] while the pathogenic potential of the European filovirus Lloviu virus (LLOV, now termed *Cuevavirus lloviuense*) [6] is incompletely understood although there is evidence that the virus might be apathogenic [7–9]. Highly pathogenic African filoviruses are considered priority pathogens by WHO and the development of vaccines and antivirals are important tasks and should be based on a thorough understanding of virus-host interactions [10].

The filovirus glycoprotein is the only viral protein that is incorporated into the viral envelope and mediates viral entry into target cells, in particular macrophages and dendritic cells [11–14]. The glycoprotein harbors a surface unit, GP1, that facilitates binding to cellular receptors while the transmembrane unit, GP2, mediates fusion of the viral membrane with the limiting membrane of late endosomes (Fig 1), which is required for release of the viral genetic information into the host cell cytoplasm [15, 16]. The GP1 surface unit engages several cellular factors for attachment to the surface of target cells in a relatively unspecific fashion while it binds with high specificity to its cognate receptor Niemann-Pick disease, type C1 (NPC1) within host cell endosomes [7, 17–21]. For this, the glycoprotein must be processed by the endosomal cysteine proteases Cathepesin B/L during viral entry into target cells [22]. Filovirus glycoproteins are also proteolytically processed upon synthesis in infected cells. Processing occurs at a motif recognized by the proprotein convertase furin [23] that is located at the interface between GP1 and GP2 (Fig 1) and furin is believed to be responsible for GP cleavage [24, 25]. In contrast, the role of proprotein convertases other than furin [23] in GP processing is less well understood. Finally, it is noteworthy that although a furin motif is present in all filovirus glycoproteins, an intact furin motif is dispensable for EBOV infection and spread in cell lines and non-human primates [26–28].

**Fig 1.**
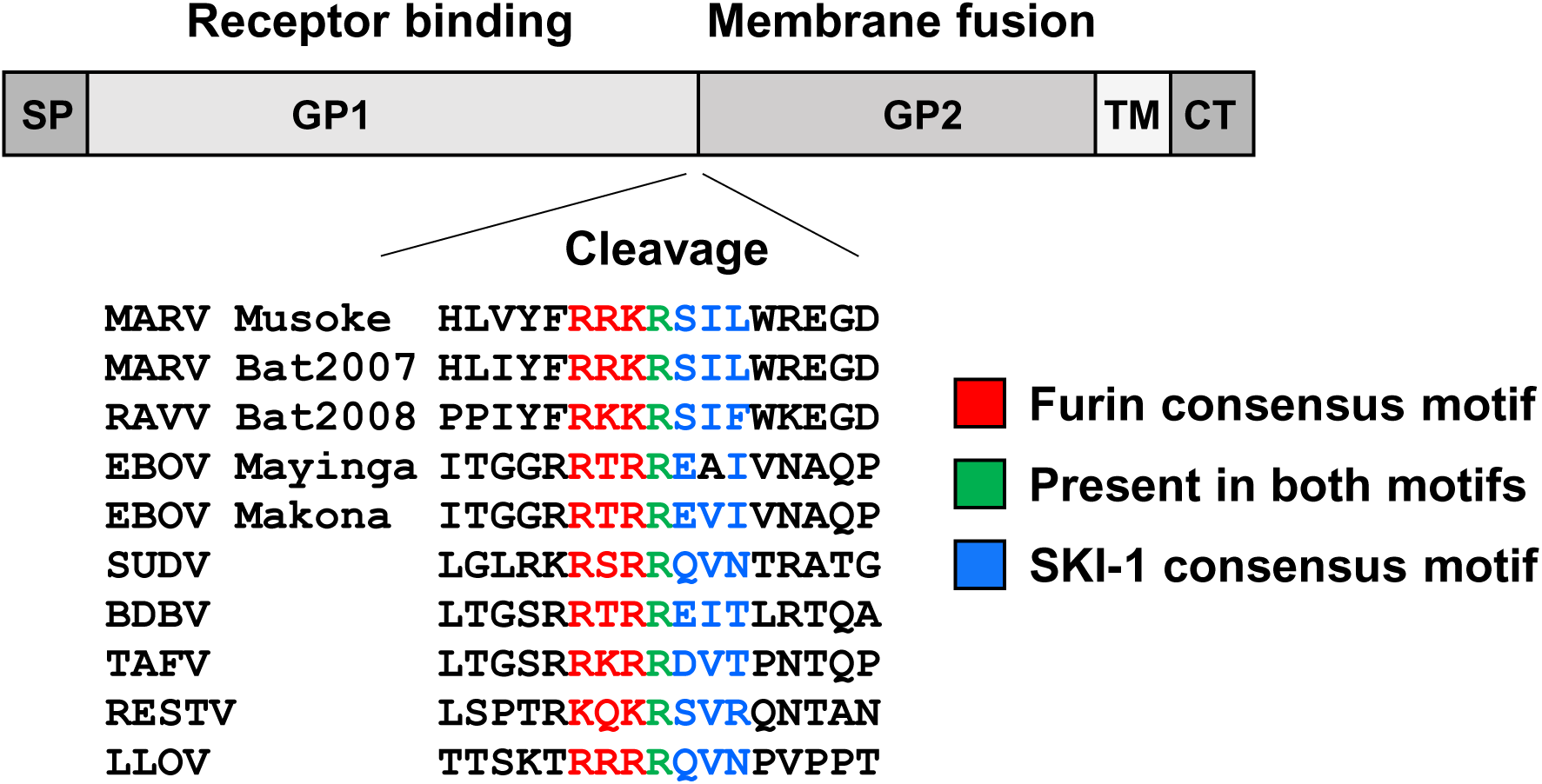
Domain organization and proprotein convertase cleavage motif in filovirus glycoproteins. Schematic overview of the domain organization of filovirus glycoproteins. The consensus motifs for furin (red) and SKI-1 (blue) as well as an arginine residue (R) that is part of both motifs are indicated. SP, signal peptide; GP1, surface unit; GP2, transmembrane unit; TM, transmembrane domain; CT, cytoplasmic tail.

The filovirus glycoproteins are expressed in the constitutive secretory pathway. Expression requires import of the nascent GP polypeptide chain into the endoplasmic reticulum (ER), which is believed to depend on recognition of an N-terminal signal peptide by the signal recognition particle (SRP) and insertion of the polypeptide into the Sec61 translocon [29]. Although many cellular proteins are imported into the ER via the same route, Zhang and colleagues showed that Sec61β knock-down (KO) was compatible with cell viability and reduced flavivirus replication [30]. Further, it has been demonstrated that the Sec61 channel can be blocked by apratoxins [31] and that Apratoxin S4 can target the translocon without compromising cellular viability and can exert antiviral activity [32, 33]. However, the role of Sec61 in filovirus infection has not been investigated, despite one report indicating that VP24 interacts with the Sec61 subunit Sec61α [34], and it is unknown whether blockade of Sec61 inhibits filovirus infection.

Here, we show that Sec61β is required for cleavage of MARV-GP but not EBOV-GP and that an intact furin motif is indispensable for efficient cell entry mediated by Marburgvirus- but not Ebola- or Cuevavirus glycoproteins. Further, we show that the proprotein convertase SKI-1 [35–37] can process MARV- but not EBOV-GP and provide evidence that SKI-1 activity is compromised in the absence of Sec61β. In addition, we show that MARV-GP N-glycosylation is altered upon Sec61β KO and identify an N-glycosylation signal that is required for MARV-GP cleavage. Finally, we demonstrate that Apratoxin S4 inhibits EBOV and MARV infection.

## Results

### Generation of *SEC61B*-KO cells

To assess the role of Sec61β in EBOV- and MARV-GP expression, Sec61β-knockout (KO) cells were generated employing CRISPR/Cas9 and single-cell cloning. Immunoblot analyses revealed that the single cell clones number 7, 9 and 10 showed no Sec61β expression while Sec61β was readily detected in WT control cells (Fig 2A). To determine a potential effect of *SEC61B*-KO on cell growth, a CFSE-based assay was employed, which relies on successive reduction of CFSE staining due to cell division (Fig 2B). The assay revealed that the growth rate of WT and *SEC61B*- KO cell lines was roughly comparable, although it should be noted that all three *SEC61B*-KO cell lines exhibited slightly higher peak fluorescence intensity after three days as compared to WT cells. Further, phase contrast microscopy images showed no appreciable differences between the morphology of WT and KO cell lines (Fig 2C). Finally, clone 9 and 10 but not clone 7 cells were less adherent as compared to WT cells (not shown) for at present unknown reason. Therefore, clone 7 cells were chosen for all further experiments. In sum, we found that *SEC61B*-KO is compatible with robust cell growth and has little if any impact on cell morphology but can reduce cell adherence.

**Fig 2.**
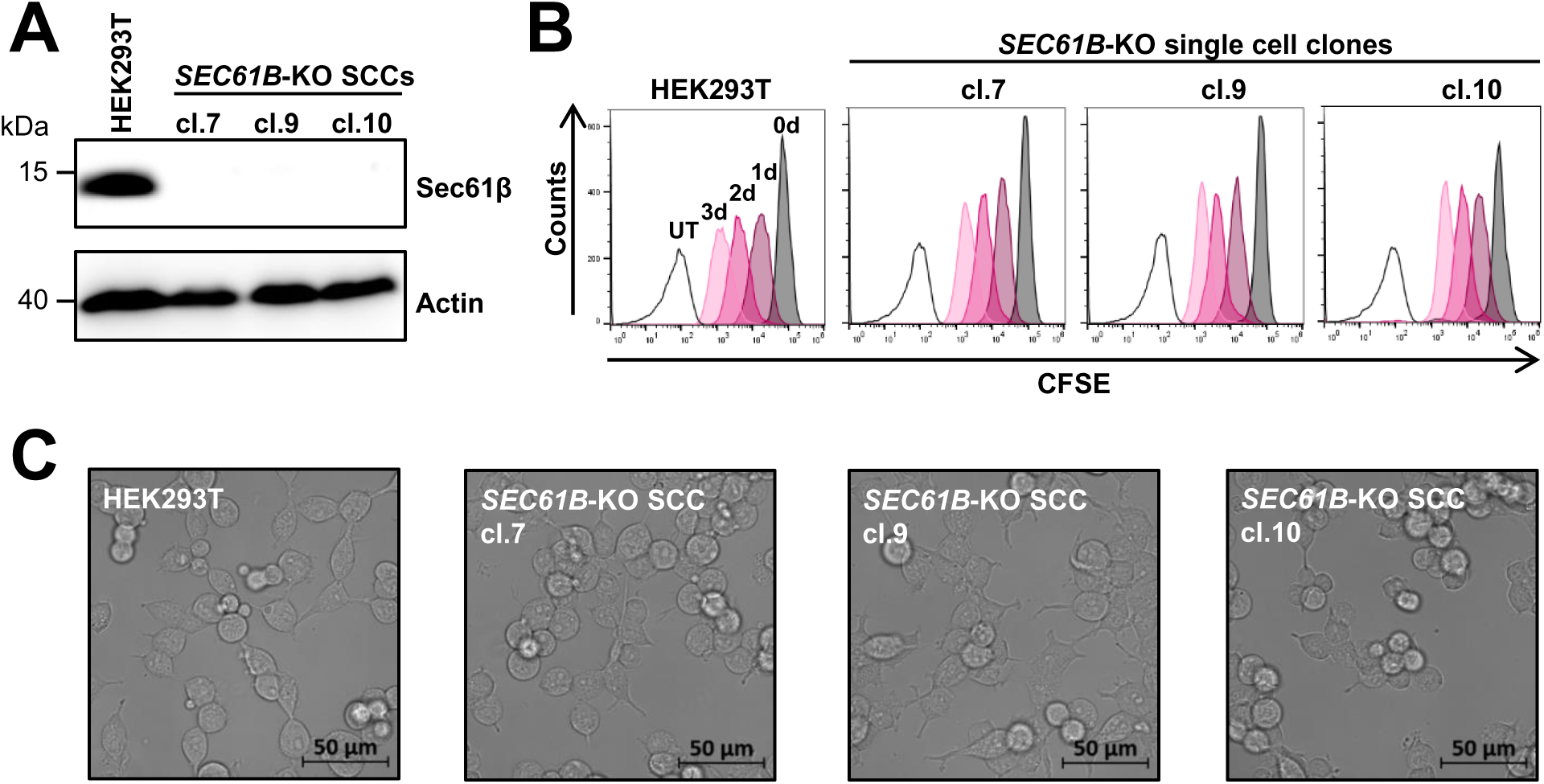
Generation of Sec61β knock-out cells. (**A**) *SEC61B*-knockout (KO) single cell clones were generated from HEK293T cells using CRISPR/Cas9. Sec61β expression in lysates of the indicated *SEC61B*-KO single cell clones was analyzed by immunoblot using polyclonal anti-Sec61β antibody. Expression of β-actin was analyzed as loading control. Results were confirmed in two additional experiments. (**B**) The indicated *SEC61B*-KO cell lines were stained with CFSE and analyzed via flow cytometry at day 0, 1, 2 and 3. The results of a single experiment are shown and were confirmed in two separate experiments. (**C**) Phase contrast microscopy of Sec61β single cell clones. Similar results were obtained in one additional experiment.

### Sec61β is required for MARV-GP but not EBOV-GP cleavage

Next, we investigated whether Sec61β affects expression of EBOV-GP (Makona) and MARV-GP (Musoke). For this, WT and *SEC61B*-KO cells were transfected with plasmids coding for EBOV- and MARV-GP with C-terminal V5-tag and analyzed by immunoblot with anti-V5-antibody. A plasmid encoding the glycoprotein of vesicular stomatitis virus (VSV-G) was included as control. Expression of VSV-G was not affected by *SEC61B*-KO and the protein was not cleaved (Fig 3A), as expected [38]. Similarly, EBOV-GP expression was robust in WT and *SEC61B*-KOcells and GP was cleaved in both cell lines. In contrast, MARV-GP cleavage was markedly reduced in *SEC61B*- KO cells, with only a faint, diffuse GP2 band being detectable that migrated slightly slower in the gel than GP2 in WT cells (Fig 3A). This effect was partially rescued by directed expression of Sec61β, which increased MARV-GP expression in WT cells and allowed for GP cleavage in *SEC61B*-KO cells, although cleavage efficiency was reduced as compared to WT cells (Fig 3B). Thus, expression of Sec61β is needed for efficient MARV-GP but not EBOV-GP cleavage.

**Fig 3.**
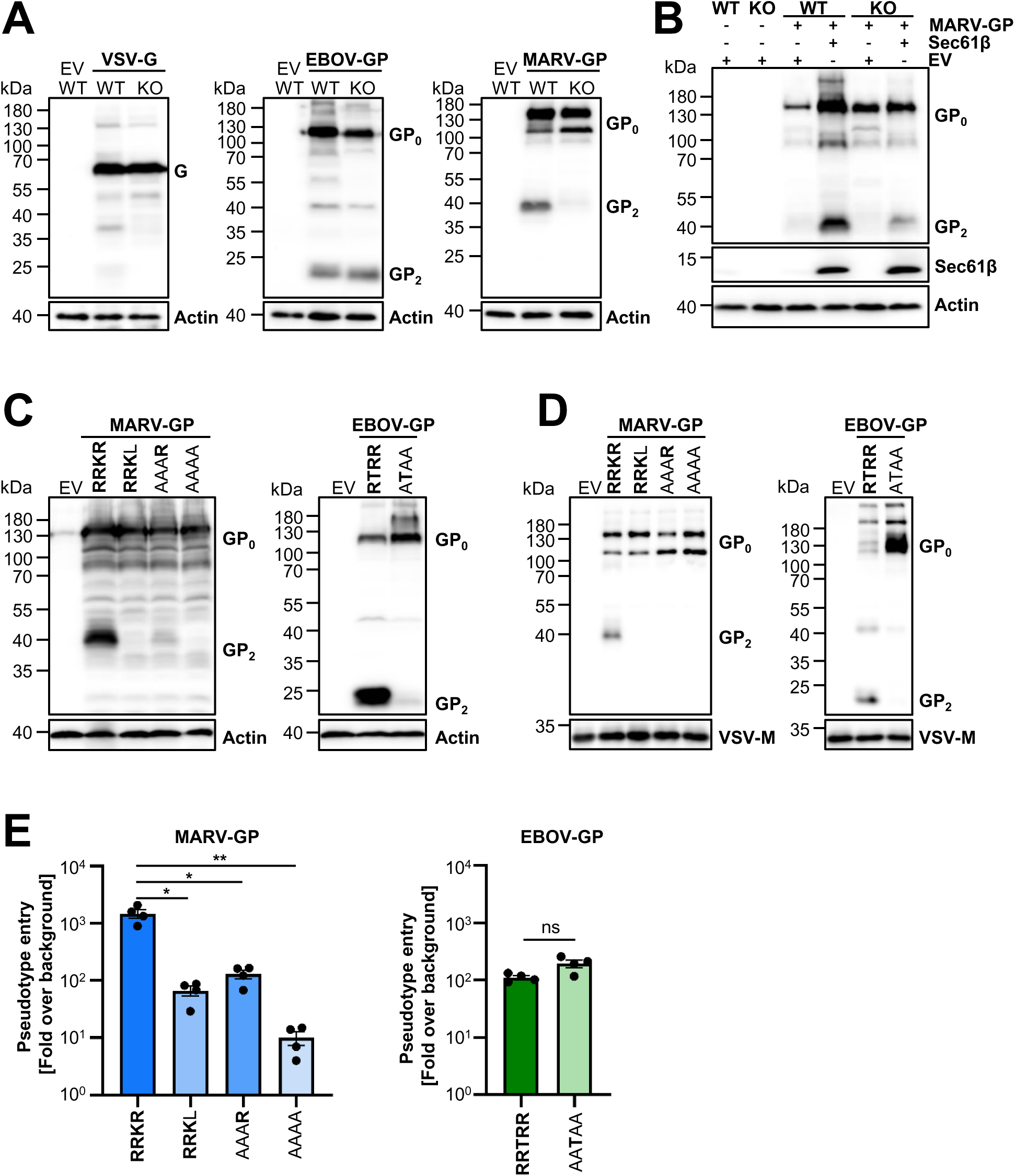
Cleavage of MARV-GP depends on Sec61β and is required for infectivity. (**A**) WT and *SEC61B*-KO cells were transfected with plasmids expressing C-terminal V5-tagged GPs of VSV, EBOV (Makona) and MARV (Musoke) or empty vector (EV) as control. Glycoprotein expression was detected by immunoblot using a mouse monoclonal anti-V5 antibody. Detection of β-actin served as a loading control. Results were confirmed by one to three additional experiments. (**B**) The experiment was conducted as for panel A but the effect of coexpression of Sec61β on MARV-GP expression and cleavage was examined. The results of a representative immunoblot are shown and were confirmed in two separate experiments. (**C-D**) The expression of MARV-GP and EBOV-GP with the indicated mutations in the furin cleavage sites was analyzed in cell lysates (C) and pseudotyped VSV particles (D). The expression of β-actin (cell lysates) and VSV-M (particles) served as loading control. Results were confirmed in two (EBOV-GP) or three (MARV-GP) additional experiments. (**E**) HEK293T cells were inoculated with VSV particles pseudotyped with the indicated glycoproteins. Luciferase activity in cell lysates were quantified at 18-20 h post inoculation. Luminescence signals were normalized to background (empty vector control). Shown is the mean ± SEM of four separate experiments each conducted with four technical replicates. Statistical significances were determined using unpaired, two-tailed t-test with Welch‘s correction (p > 0.05, not significant [ns]; p ≤ 0.05, *; p ≤ 0.01, **)

### An intact furin motif is required for cell entry driven by MARV-GP but not EBOV-GP glycoproteins

To assess whether reduced MARV-GP cleavage might compromise MARV-GP-driven entry, we mutated the furin cleavage site located between the surface unit GP1 and the transmembrane unit GP2. As control, we also mutated the furin motif in EBOV-GP since the cleavage site is known to be dispensable for EBOV-GP expression and function [26–28]. Specifically, we exchanged the furin motif in MARV-GP from RRKR to RRKL, considering that the R to L exchange was previously shown to abrogate MARV-GP cleavage [25]. In addition, the RRKR motif was changed to AAAR and AAAA while the furin cleavage motif in EBOV-GP, RRTRR, was changed to AATAA, as reported before [26–28, 39]. All mutant GPs were robustly expressed in transfected cells (Fig 3C) and incorporated into VSV particles (Fig 3D). Mutation of the MARV-GP furin cleavage motif, RRKR, to AAAR reduced GP cleavage, although a cleavage product was still readily detectable, while mutation to RRKL largely and mutation to AAAA completely abrogated cleavage (Fig 3C, D). Similarly, mutation AATAA in EBOV-GP abrogated GP cleavage (Fig 3C, D). Finally and notably, mutations in the MARV-GP furin motif reduced GP-driven cell entry of VSV pseudoparticles, with a marked correspondence between cleavage and entry efficiency, while mutation of the furin motif in EBOV-GP did not diminish entry efficiency (**Fig 3E**), as expected. In sum, an intact furin motif is required for cell entry driven by MARV-GP but not EBOV-GP.

### An intact furin motif is dispensable for cell entry driven by the glycoproteins of Ebola- and Cuevaviruses but not Marburgviruses

The analyses of GP expression and GP-mediated cleavage were so far conducted with MARV-GP Musoke and EBOV-GP Makona. Therefore, we expanded our studies to the glycoproteins of EBOV (Mayinga), Sudan virus (SUDV), Tai Forrest virus (TAFV), Reston virus (RESTV), Bundibugyo virus (BDBV), Lloviu virus (LLOV), bat-derived MARV (Bat2007) and Ravn virus (RAVV, Bat2008). Mutation of the furin cleavage motifs in these GPs largely abrogated cleavage (Fig 4A) but had differential effects on GP-driven entry. Thus, particles bearing EBOV-GP (Mayinga, and Makona), SUDV-GP, TAFV-GP, BDBV-GP, RESTV-GP and LLOV-GP with mutated furin motif showed comparable or increased infectivity as compared to their counterparts harboring intact furin motifs (Fig 4B). Further, mutation of the furin motif in MARV-GP (Bat2007) and RAVV-GP slightly but significantly reduced entry (Fig 4B). Finally, mutation of the furin motif in MARV-GP Musoke markedly reduced entry, as expected. These results suggest that an intact furin motif might be dispensable for Ebola (EBOV, SUDV, BDBV, TAFV, RESTV) and Cuevaviruses (LLOV) while the same motif might be of moderate (RAVV, MARV Bat2007) or high (MARV Musoke) importance for Marburg viruses.

**Figure 4.**
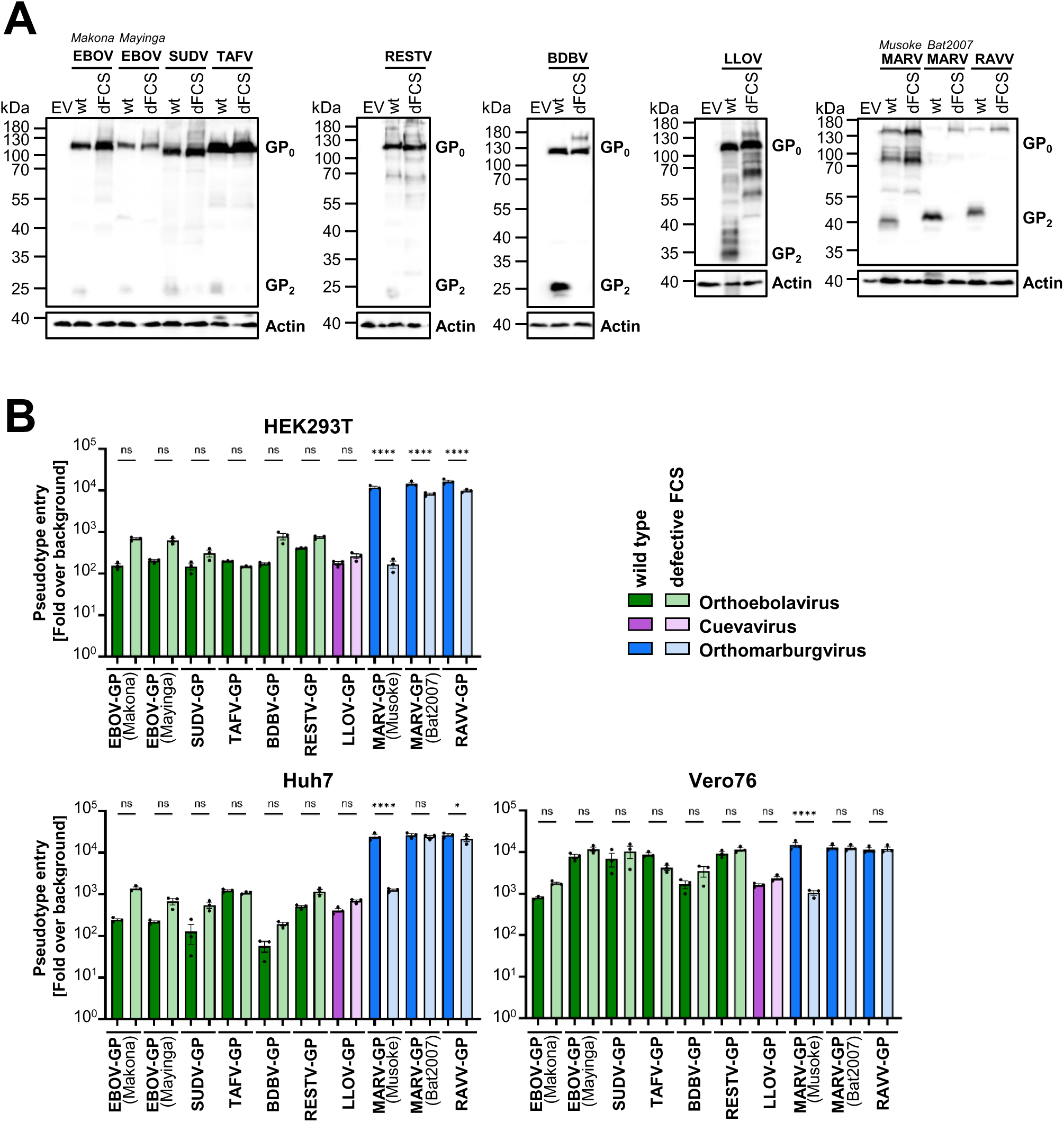
An intact furin motif is required for infectivity of particles bearing the glycoproteins of Marburg- but not Ebola- or Cueavaviruses. (**A**) The indicated viral glycoproteins equipped with a C-terminal V5-tag harboring either an intact (wt) furin cleavage site (FCS) or a defective FCS (dFCS) were expressed in HEK293T WT cells. Expression was analyzed by immunoblot using a mouse monoclonal anti-V5 antibody. Detection of β-actin served as a loading control. Results were confirmed in one to two additional experiments. (**B**) VSV pseudoparticles bearing the indicated GPs with either WT FCS or dFCS were used to infect HEK293T, Huh7 and Vero76 cells. After 18-20 h, the luciferase activity in cell lysates was determined. Shown is the mean ± SEM of three individual experiments with technical quadruplicates. Statistical significance was determined using one-way ANOVA with Šidák correction (p > 0.05, not significant [ns]; p ≤ 0.05, *; p ≤ 0.01, **).

### Directed expression of furin in *SEC61B*-KO cells does not rescue MARV-GP cleavage

Next, we investigated whether the largely absent cleavage of MARV-GP in *SEC61B*-KO cells could be rescued by directed expression of furin. For control purposes, the effect of directed furin expression on EBOV-GP cleavage was analyzed in parallel. Endogenous and exogenous (i.e. overexpressed) furin were readily detectable in transfected WT and *SEC61B*-KO cells (Fig 5A). In WT cells, directed furin expression markedly increased MARV-GP expression but not cleavage while directed furin expression in *SEC61B*-KO cells neither modulated MARV-GP expression nor cleavage (Fig 5A, left panel). In the context of EBOV-GP expression, directed furin expression in WT cells markedly increased EBOV-GP cleavage but not expression while no effect of directed furin expression was observed in *SEC61B*-KO cells (Fig 5A, right panel). These results suggest that endogenous furin levels might limit cleavage of EBOV-GP but not MARV-GP in WT cells. Further, based on the absence of MARV-GP cleavage in *SEC61B*-KO cells overexpressing furin and the lack of enhancement of EBOV-GP cleavage in these cells, the results indicate that Sec61β is required for the ability of furin to cleave GP.

**Figure 5.**
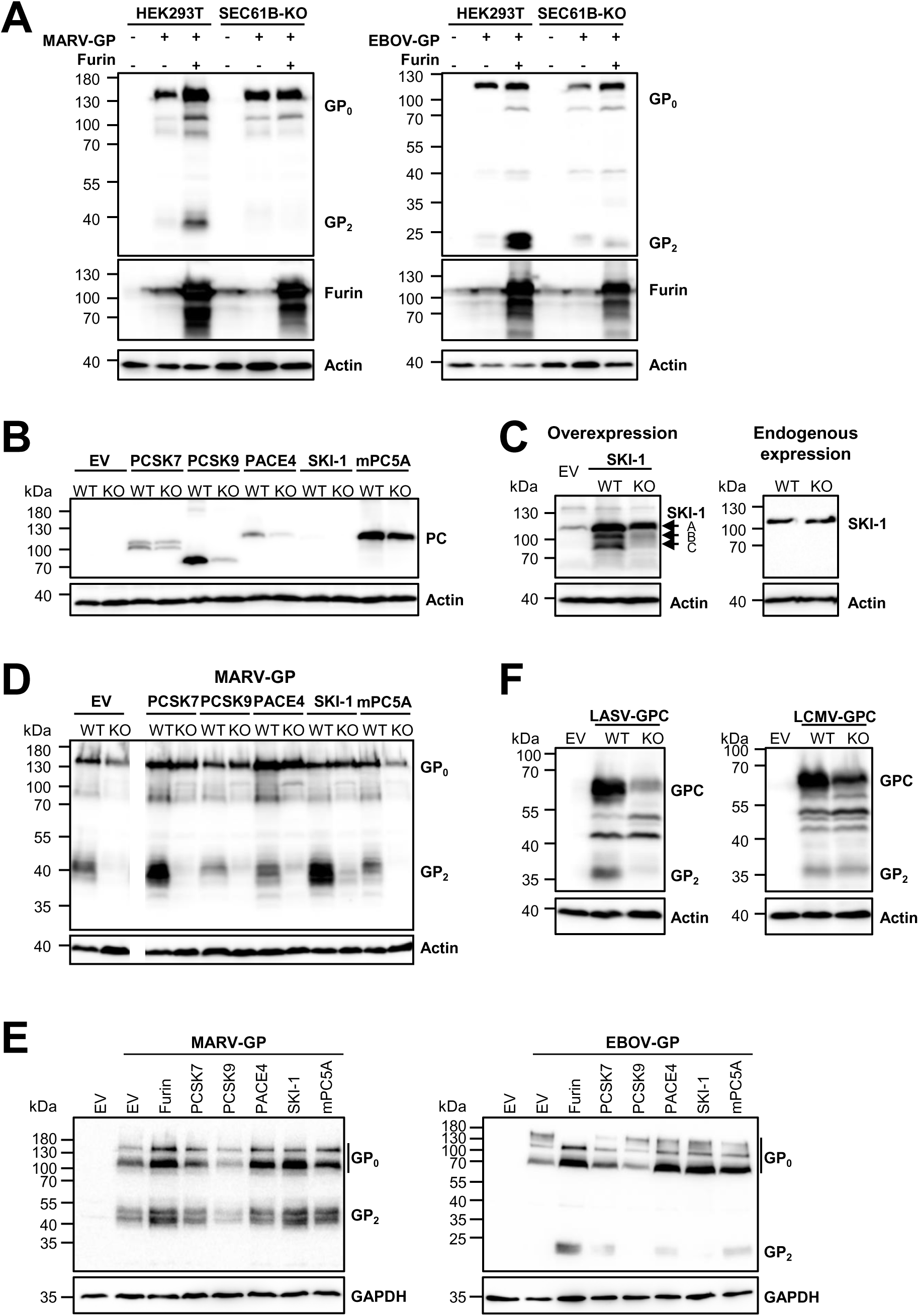
MARV-GP is cleaved by SKI-1 and SKI-1-mediated cleavage is compromised in *SEC61B*-KO cells. (**A**) MARV-GP and EBOV-GP with a C-terminal V5-tag were coexpressed with furin or empty vector (EV) in WT or *SEC61B*-KO cells and expression of GP analyzed by immunoblot using anti- V5 antibody. In parallel, expression of furin was determined using a rabbit polyclonal-anti furin antibody and expression of β-actin was analyzed as loading control. The results were confirmed in two additional experiments. (**B**) Expression of cMyc-tagged PCSK7, PCSK9, PACE4, SKI-1 and mPC5A in WT and *SEC61B*-KO cells was assessed by immunoblot using anti-cMyc antibody. Detection of β-actin served as a loading control. Similar results were obtained in six additional experiments. (**C**) Directed expression of cMyc-tagged SKI-1 in WT and *SEC61B*-KO cells was analyzed by immunoblot using a polyclonal SKI-1 specific antibody. Endogenous expression of SKI-1 was detected using a monoclonal antibody recognizing the N-terminal domain of SKI-1. Results were confirmed in two (overexpression) or one (endogenous expression) additional experiments. (**D**) The indicated cMyc-tagged proprotein convertases (PCs) PCSK7, PCSK9, PACE4, SKI-1 and mPC5A were coexpressed with MARV-GP harboring a V5-antigenic tag and GP expression analyzed by immunoblot. Transfection of empty vector (EV) served as specificity control, detection of β-actin served as a loading control. Similar results were obtained in two separate experiments. (**E**) The experiment was carried out as described for panel (D) but LoVo cells, which are defective for furin, were used and EBOV-GP was included in the experiment. Similar results were obtained in two separate experiments. (**F**) LASV-GPC or LCMV-GPC with a C-terminal V5-tag were expressed in WT and *SEC61B*-KOcells and expression analyzed by immunoblot. Transfection of empty vector (EV) served as specificity control, detection of β-actin served as loading control. Similar results were obtained in two separate experiments.

### SKI-1 cleaves MARV-GP in WT cells and directed expression of SKI-1 in *SEC61B*-KO cells moderately rescues MARV-GP cleavage

We next investigated whether the directed expression of proprotein convertases other than furin allows for MARV-GP cleavage in *SEC61B*-KO cells. For this, proteases bearing a cMyc-tag at the C-terminus were used. Directed expression of PCSK7, PCSK9, PACE4 and murine (m) PC5A in WT cells was readily detectable and more efficient than that observed in *SEC61B*-KO cells (Fig 5B). Directed expression of SKI-1 in WT cells was barely detectable and we failed to detect expression in *SEC61B*-KO cells, using the anti-cMyc antibody (Fig 5B). In contrast, directed SKI-1 expression in WT and *SEC61B*-KO cells was readily detectable using an anti-SKI-1 antibody and revealed that processing of SKI-1 into its active 96/102 kDa forms was less efficient in *SEC61B*-KO as compared to WT cells (Fig 5C). Endogenous SKI-1 expression was also detected using an antibody specific for the inactive form and no differences in SKI-1 expression levels in WT and *SEC61B*-KO cells were noted (Fig 5C). Thus, all proprotein convertases tested were expressed in both WT and *SEC61B*-KO cells but differences in expression efficiency were noted.

Coexpression of proprotein convertases with MARV-GP revealed that PCSK7 and SKI-1 markedly increased MARV-GP cleavage in WT cells while this effect was not observed for cells coexpressing the other proprotein convertases or empty vector (Fig 5D). In contrast, none of the proprotein convertases tested efficiently rescued MARV-GP cleavage in *SEC61B*-KO cells although inefficient rescue was consistently observed upon SKI-1 overexpression (Fig 5D). We next sought confirm these results in an independent cell system and to extend our analysis to EBOV-GP. For this, we employed LoVo cells, which do not express enzymatically active furin [40]. Notably, MARV-GP was appreciably cleaved in LoVo control cells transfected with empty vector (EV) and cleavage was augmented upon directed expression of furin and SKI-1 (Fig 5E, left panel). In contrast, EBOV-GP was uncleaved in LoVo control cells and only directed expression of furin resulted in robust GP cleavage, although inefficient GP cleavage was detected upon directed expression of PCSK7, PACE4 and mPC5A (Fig 5E, right panel). Collectively, these results demonstrate that MARV-GP but not EBOV-GP can employ SKI-1 in addition to furin for GP cleavage and suggest that the ability of endogenous SKI-1 to cleave MARV-GP in *SEC61B*-KO cells might be compromised.

### The Lassa virus glycoprotein, a known SKI-1 substrate, is not efficiently processed in *SEC61B*-KO cells

In order to confirm that the ability of SKI-1 to cleave viral glycoproteins is indeed compromised in *SEC61B*-KO cells, we investigated cleavage of the Lassa virus (LASV) and Lymphocytic choriomeningitis virus (LCMV) glycoproteins (GPC), two well-characterized substrates of SKI-1 [41–43]. Both proteins were cleaved in WT cells, as expected, but only LCMV-GPC was efficiently cleaved in *SEC61B*-KO cells (Fig 5F). Notably, SKI-1-mediated cleavage of LCMV-GP occurs in the late Golgi [41, 44] while cleavage of LASV-GPC proceeds in the ER/cis-Golgi [42, 43], suggesting that the ability of SKI-1 to process viral glycoproteins in the ER/cis-Golgi might be reduced in *SEC61B*-KO cells.

### SKI-1 inhibitors block infection by recombinant vesicular stomatitis virus chimeras encoding EBOV-GP and MARV-GP

To further investigate the potential role of SKI-1 in MARV infection, we used two SKI-1 inhibitors, PF-429242 [45] and MI-2701 (Supplementary figure S1) as well as recombinant vesicular stomatitis virus encoding MARV-GP instead of the VSV glycoprotein (rVSV-MARV-GP). As controls, recombinant VSV (rVSV) as well as rVSV-LASV-GPC and rVSV-EBOV-GP were also examined. The antiviral activity of Decanoyl-RVKR-CMK, a furin inhibitor, FI, [46], was analyzed in parallel. The furin inhibitor did not inhibit infection with rVSV and rVSV-LASV-GPC and only moderately inhibited rVSV-MARV-GP and rVSV-EBOV-GP (Fig 6A). The SKI-1 inhibitors were largely inactive against rVSV but potently blocked rVSV-LASV-GP, as expected [47–49]. Notably, both SKI-1 inhibitors also potently blocked rVSV-MARV-GP and rVSV-EBOV-GP infection (Fig 6A). Finally, combining the furin inhibitor with the SKI-1 inhibitor PF-429242 slightly inhibited rVSV, potentially due to unspecific effects, and blocked rVSV-LASV-GPC, rVSV-MARV-GP and rVSV-EBOV-GP with similar efficiency as PF-429242 alone. Thus, the activity of endogenous SKI-1 is required for EBOV-GP- and MARV-GP-dependent infection, at least in the HEK293T cells analyzed here.

**Figure 6.**
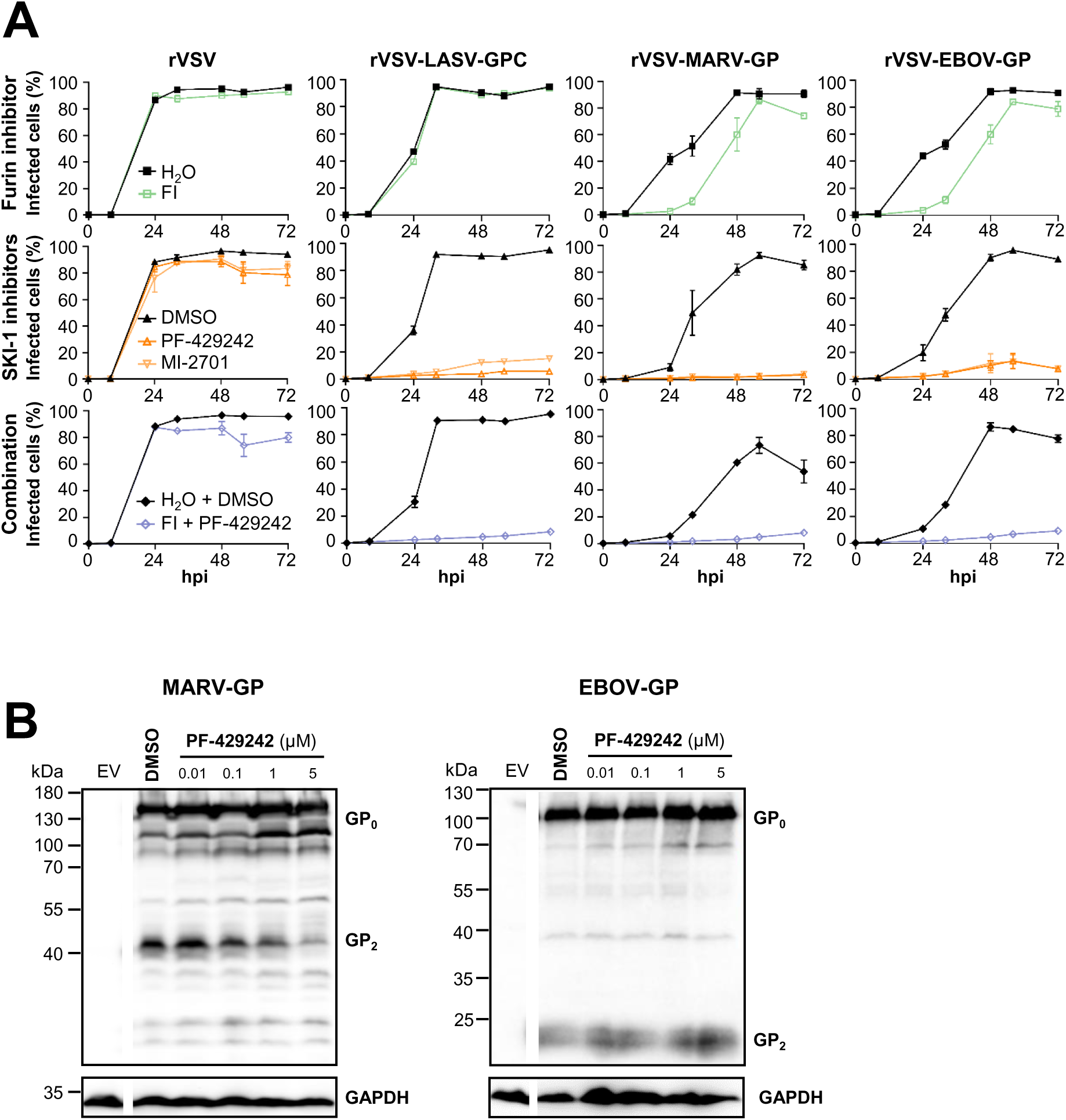
SKI-1 inhibitors block infection by recombinant vesicular stomatitis virus chimeras encoding EBOV-GP and MARV-GP. (**A**) HEK293T WT cells were infected with rVSV, rVSV-LASV-GPC, rVSV-MARV-GP and rVSV-EBOV-GP at 0.01 MOI. After 1 h, inoculum was replaced with medium containing either furin inhibitor (Decanoyl-RVKR-CMK, 25 µM; H_2_O as control) or SKI-1 inhibitors (PF-429242, 5 µM or MI-2701, 1 µM; DMSO as control) or an inhibitor combination (25 µM decanoyl-RVKR- CMK and 5 µM PF-429242; H_2_O and DMSO as control). Cells were harvested at the indicated time points, fixed and analyzed for GFP^+^ cells via flow cytometry. Shown is the mean (+/- SEM) percentage of infected cells from three individual experiments. (**B**) HEK293T WT cells transfected to express MARV-GP or EBOV-GP with V5-antigenic tag were treated with the SKI-1 inhibitor PF-429242 and GP expression was analyzed via immunoblot with a V5-specific antibody. Visualization of GAPDH served a loading control. Results were confirmed in two independent experiments.

A previous study demonstrated that processing of N-acetylglucosamine-1-phosphate transferase α and β subunit (GNPTAB) by SKI-1 is required for GNTAB activity [50], which in turn is essential for robust expression of cathepsin B, a cellular protease that cleaves filovirus GPs and supports filovirus entry into cells [22, 48]. Therefore, we investigated whether the inhibition of GP cleavage might have contributed to the antiviral activity of the SKI-1 inhibitors. For this, we examined inhibition of MARV-GP and EBOV-GP cleavage by PF-429242. Treatment with PF- 429242 markedly and dose-dependently inhibited MARV-GP cleavage but had only a minor effect on EBOV-GP cleavage (Fig 6B), indicating that endogenous SKI-1 can process MARV-GP and that inhibition of processing might have contributed to the anti-rVSV-MARV-GP activity of PF- 429242.

### N-glycosylation of MARV-GP in *SEC61B*-KO cells is heterogeneous and mutation N564D, which destroys a sequon, reduces MARV-GP cleavage

Next, we investigated whether determinants in MARV-GP might also contribute to the inefficient cleavage in *SEC61B*-KO cells. For this, we focused on N-glycosylation since it was previously shown that mutation of sequons in GP2 of EBOV-GP impacts GP cleavage and cell entry efficiency [51–53]. Additionally, an important role of N-glycosylation in MARV-GP transport through the secretory pathway has been shown using tunicamycin [54]. We first asked whether the faint, diffuse GP2 band observed in *SEC61B*-KO cells expressing MARV-GP might represent different glycoforms of GP. Indeed, digest of cell lysates with PNGaseF, which removes all N-linked glycans, resulted in sharp GP2 bands in both WT and *SEC61B*-KO cells and both bands exhibited the same molecular weight, although the GP2 band detected in KO cells was less prominent than its counterpart in WT cells (Fig 7A, left panel). In contrast, no differences in GP2 bands in the absence or presence of PNGaseF treatment was detected for WT and *SEC61B*-KO cells expressing EBOV-GP (Fig 7A, right panel), as expected. These results suggest that MARV-GP N-glycosylation in *SEC61B*-KO cells is heterogeneous, raising the possibility that reduced MARV-GP cleavage might in part stem from inefficient or absent usage of certain sequons within GP2.

**Figure 7.**
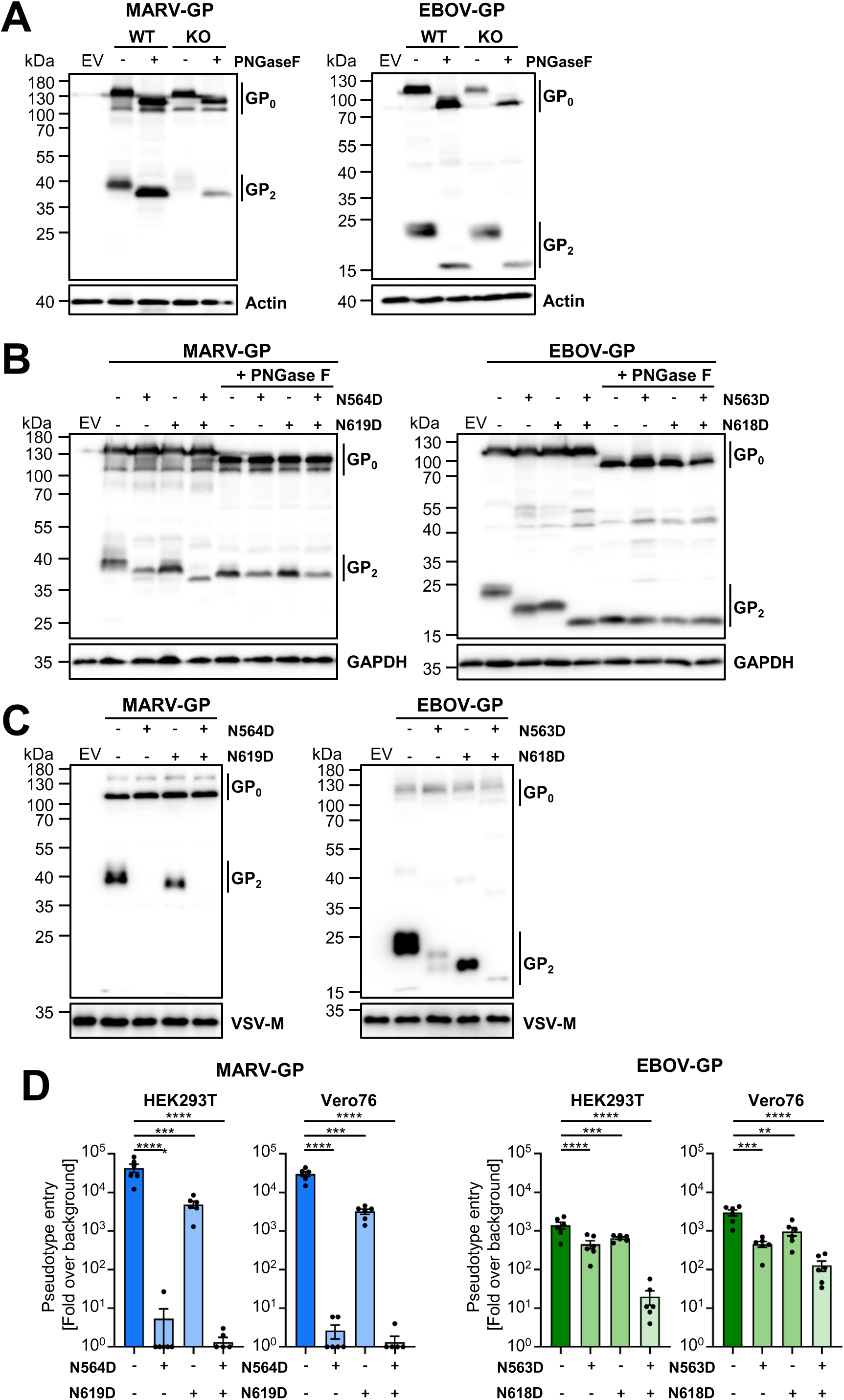
*SEC61B*-KO interferes with MARV-GP N-glycosylation and mutation N564D, which destroys a sequon, abrogates MARV-GP cleavage. (**A**) MARV-GP and EBOV-GP with C-terminal V5-tag were expressed in WT and *SEC61B*-KO cells, cell lysates treated with PNGase F or control treated and GP expression in cell lysates analyzed by immunoblot using a mouse monoclonal anti-V5 antibody. Cells transfected with empty vector (EV) served as specificity control. Detection of β-actin served as loading control. The results were confirmed in two separate experiments. (**B**) The experiment was performed as described for panel A but the indicated MARV-GP and EBOV-GP sequon mutants were analyzed. Detection of GAPDH served as loading control. Results were confirmed in three (for MARV-GP) or one (for EBOV-GP) separate experiments. (**C**) Incorporation of the indicated MARV-GP and EBOV-GP sequon mutants with C-terminal V5-tag in VSV particles was analyzed by immunoblot using a V5- specific antibody. Detection of VSV-M served as loading control. Similar results were obtained in three additional experiments. (**D**) Equal volumes of VSV particles pseudotyped with the indicated glycoproteins were inoculated onto HEK293T and Vero76 cells. After 18-20 h luciferase activity in cell lysates was measured. Luminescence signals were normalized to background (empty vector control). Shown are the mean values ± SEM of six independent experiments with technical quadruplicates. Statistical significances were tested using One-Way ANOVA with Dunnett correction (p > 0.05, not significant [ns]; p ≤ 0.05, *; p ≤ 0.01, **; p ≤ 0.001, ***; p ≤ 0.0001, ****).

The sequons compromising N563D and N618D were previously shown to be required for particle incorporation of cleaved EBOV-GP and combining both mutations markedly reduced infectivity [51–53]. Therefore, we examined the effect of mutation of these residues and their counterparts in MARV-GP (N564D and N619D) on GP glycosylation, cleavage and ability to drive cell entry. Expression of these mutants in WT cells followed by mock treatment or treatment with PNGaseF revealed that both N564 and N619 in MARV-GP were N-glycosylated and that mutation N564D reduced MARV-GP cleavage (Fig 7B, left panel). Similarly, both N563 and N618 were N-glycosylated in EBOV-GP but neither N563D nor N618D interfered with GP cleavage (Fig 7B, right panel). More pronounced effects were observed when GPs were examined that were incorporated into VSV particles and thus had completed their passage through the constitutive secretory pathway. Thus, mutation N564D abrogated incorporation of cleaved MARV-GP into particles and the corresponding mutation in EBOV-GP, N563D, markedly reduced particle incorporation of cleaved GP (Fig 7C). Further, mutation N619D in MARV-GP and N618D in EBOV-GP moderately reduced the levels of particle-associated cleaved GP, and mutating both sequons further reduced EBOV-GP cleavage (Fig 7C). In keeping with these findings, mutation N564D not only abrogated particle incorporation of cleaved MARV-GP but also reduced particle infectivity close to background levels while mutation N619D had only a moderate effect on infectivity (Fig 7D, left panel). In contrast, mutations of N564D and N619D in EBOV-GP had only a moderate effect on infectivity (Fig 7D, right panel), in agreement with the notion that cleavage is dispensable for GP-driven entry. Finally, combining N563D and N618D in EBOV-GP markedly reduced infectivity but this effect might have been due to reduced particle incorporation or other factors. In sum, the absence of Sec61β interferes with N-glycosylation of MARV-GP and mutation of the sequon comprising N564 inhibits MARV-GP cleavage.

### Blockade of Sec61 by Apratoxin S4 inhibits EBOV and MARV infection

We finally explored whether Sec61β or the entire Sec61 translocon might constitute targets for antiviral intervention. Inoculation of WT and *SEC61B*-KO cells with rVSV, rVSV-EBOV-GP and rVSV-MARV-GP followed by TCID_50_ quantification in culture supernatants revealed comparable infection of both WT and *SEC61B*-KO cells with rVSV (Fig 8A). Infection of WT cells with rVSV-MARV-GP and particularly rVSV-EBOV-GP was less efficient than that measured for rVSV and *SEC61B*-KO further reduced infection, although this effect was not statistically significant (Fig 8A). The analysis of glycoprotein incorporation into single cycle particles revealed that although VSV-G was comparably incorporated into particles produced in WT and *SEC61B*-KO cells, incorporation of MARV-GP and EBOV-GP was markedly reduced and probably accounted for the reduced infection efficiency (Fig 8B). Finally, authentic MARV (Musoke) or EBOV (Mayinga) also infected *SEC61B*-KO cells with reduced efficiency as compared to WT cells but these effects were moderate and not statistically significant (Fig 8C). These findings suggest that Sec61β might be required for EBOV- and MARV-GP transport to the side of VSV budding, the plasma membrane. Furthermore, the data indicate that Sec61β might promote but is not essential for MARV and EBOV infection.

**Figure 8.**
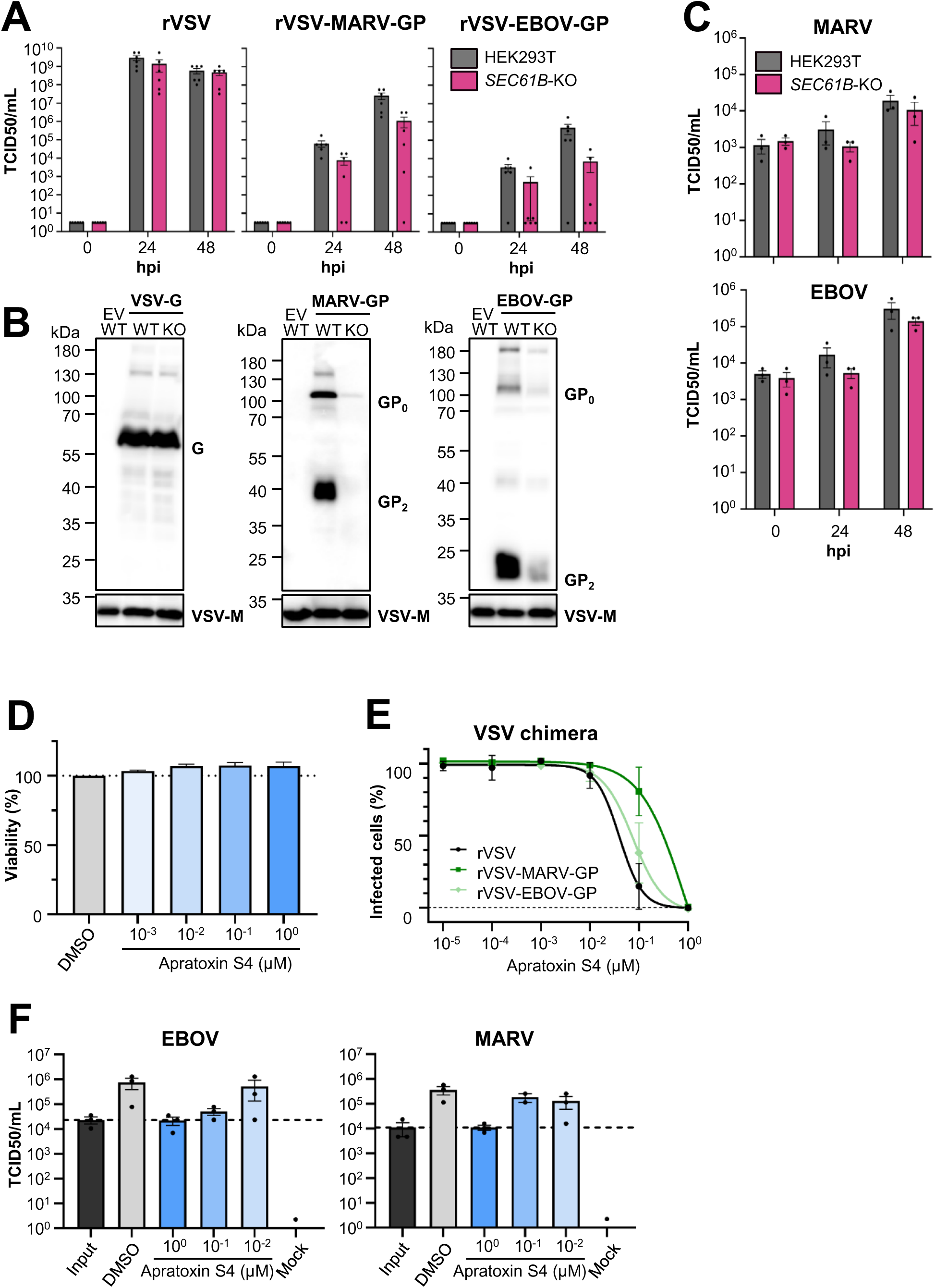
Apratoxin inhibits EBOV and MARV infection. (A) WT or *SEC61B*-KO cells were inoculated with replication-competent, chimeric rVSV encoding VSV-G, EBOV-GP (Makona strain) or MARV-GP (Musoke strain) at an MOI of 0.01. The amount of infectious viral particles in the supernatant at the indicated time points after infection was determined by TCID50. The mean ± SEM of six separate experiments is shown. (**B**) VSV particles pseudotyped with the indicated GPs bearing a V5-antigenic tag were concentrated and GP expression analyzed by immunoblot with a V5-specific antibody. VSV-M expression is shown as loading control. Results were confirmed in two additional experiments. (**C**) The experiment was carried out as described for panel A but authentic MARV (Musoke strain) and EBOV (Mayinga strain) were used. Data represent the mean ± SEM of three separate experiments. (**D**) Vero76 cells were treated with the indicated concentrations of Apratoxin S4 or DMSO. ATP levels reflecting the cell viability were measured 50 h after treatment using the CellTiter-Glo® Luminescent Cell Viability Assay from Promega. Luminescence signals were normalized to DMSO control and displayed as relative viability. Shown is the mean ± SD of two individual experiments. (**E**) Vero76 cells were treated with the indicated concentration of Apratoxin S4 or DMSO for 2 h followed by infection (MOI 0.01) with EBOV (Mayinga) and MARV (Musoke). The amount of infectious virus in culture supernatants at 48 hpi was determined by TCID50. Infectious units per mL were calculated according to the Spearman-Kärber method. The mean ± SEM of three independent experiments is presented. (**F**) Vero 76 cells were infected 2 h after treatment with 1-10^-2^ µM Apratoxin S4 or DMSO with 0.1 MOI authentic MARV (Musoke strain) and EBOV (Mayinga strain). The amount of infectious virus at 48 hpi was determined by TCID_50._ Infectious units per mL were calculated according to the Spearman Kärber method. The mean and SEM of three independent experiments is shown.

We next investigated whether blockade of the Sec61 channel rather than interfering with expression of a channel component, Sec61β, results in more pronounced antiviral activity. For this, we employed Apratoxin S4, which is a small molecule inhibitor that stabilizes the Sec61 channel in its closed conformation [55]. Apratoxin S4 concentrations up to 1 µM were not cytotoxic in Vero76 cells (Fig 8D) but reduced infection with rVSV, rVSV-EBOV-GP and rVSV-MARV-GP to background levels (Fig 8E). Furthermore, Apratoxin S4 inhibited EBOV and MARV infection of Vero76 cells in a concentration-dependent manner, with 1 µM Apratoxin exhibiting potent antiviral activity (Fig 8F). These results suggest that blockade of the Sec61 channel rather than interference with Sec61β could be a viable therapeutic strategy against filoviruses.

## Discussion

We report that cleavage of MARV-GP and incorporation of both MARV-GP and EBOV-GP into VSV particles was markedly reduced in the absence of Sec61β. Cleavage of MARV-GP but not EBOV-GP was required for infectious GP-driven entry and MARV-GP but not EBOV-GP was able to employ SKI-1 for GP cleavage. In addition, our findings suggest that SKI-1 activity is reduced and N-glycosylation of MARV-GP is altered in *SEC61B*-KO cells and that both may contribute to inefficient MARV-GP cleavage. Finally, we found that the blockade of the Sec61 translocon by Apratoxin S4 inhibits EBOV and MARV infection.

Our finding that *SEC61B*-KO is compatible with cell viability (Fig 2) is in keeping with previous studies suggesting that Sec61β might play a stabilizing and/or regulatory but not an essential role in Sec61 activity [30, 56–59]. We observed that *SEC61B*-KO can interfere with adherence of cells to plastic plates and markedly reduces transfectability with the calcium-phosphate- but not the PEI-based method (not shown) and the underlying reasons remain to be determined.

*SEC61B*-KO markedly reduced MARV- but not EBOV-GP cleavage (Fig 3), raising the question whether reduced cleavage might result in diminished GP-driven entry. We found that the integrity of the furin cleavage motif was dispensable for cell entry driven by the GPs of several members of the *Orthoebolavirus* and *Cuevavirus* genera, in keeping with expectations for EBOV [26–28, 60], while mutation of the furin motif reduced entry driven by all members of the Orthomarburgvirus genus tested, with the most pronounced effect measured for MARV-GP Musoke (Fig 4). Thus, there appears to be a general difference between Orthoebolaviruses and Cuevaviruses on the one hand and Orthomarburgviruses on the other hand regarding the reliance on the furin motif for GP activation. These findings await conformation with authentic filoviruses and the reason(s) for these differences remain to be explored. Mutagenic analyses of Orthomarburgvirus GPs might reveal which factors determine the need for an intact furin motif. However, such endeavors might be cumbersome since MARV-GP Musoke (marked need for intact furin motif) and MARV-GP Bat2007 (minor need for intact furin motif) harbor an identical furin motif but differ in 60 amino acids in the remainder of GP.

Ebola- and Marburgvirus GPs not only differed in the requirement for an intact furin motif for cell entry but also differed in the spectrum of proprotein convertases that can cleave GP. Thus, directed expression of SKI-1 and furin resulted in robust MARV-GP cleavage while only directed expression of furin but not other proprotein convertases allowed for efficient EBOV-GP cleavage (Fig 5). Cleavage of GP upon directed overexpression of proprotein convertases was not observed in *SEC61B*-KO cells, although a slight augmentation of MARV-GP cleavage was detected upon directed SKI-1 expression. Further, *SEC61B*-KO interfered with SKI-1-dependent cleavage of LASV- but not LCMV-GPC, although both proteins are well-established substrates of SKI-1 [41–43], and it is known that cleavage of LCMV-GP occurs in the late Golgi [41, 44] while cleavage of LASV-GPC proceeds in the ER/cis-Golgi [42, 43]. These results identified SKI-1 as a novel MARV-GP processing protease and showed that GP cleavage by SKI-1 and other proprotein convertases is compromised in *SEC61B*-KO cells.

The SKI-1 inhibitors PF-429242 and MI-2701 blocked rVSV-MARV-GP and rVSV-EBOV-GP infection (Fig 6), indicating that the activity of endogenous SKI-1 is essential for MARV- and EBOV-GP-driven cell entry. However, it is unclear to which extent the antiviral activity of these inhibitors stems from blockade of GP cleavage. Thus, the cellular enzyme N-acetylglucosamine-1-phosphate (GNPTAB) is activated by SKI-1 and its activity is essential for robust expression of cathepsin B [50], a cellular cysteine protease that activates diverse filovirus GPs [22, 61, 62]. Although MARV is apparently less cathepsin B-dependent as compared to other filoviruses [62, 63], *GNPTAB*-KO still markedly reduced MARV infection [50]. We found that PF-429242 inhibited MARV-GP but not EBOV-GP cleavage in transfected cells in a concentration-dependent fashion, demonstrating that endogenous SKI-1 cleaves MARV-GP and indicating that inhibition of rVSV-MARV-GP by SKI-1 inhibitors might at least partially stem from inhibition of MARV-GP cleavage.

The cleavage of MARV-GP by SKI-1 was unexpected since the consensus motif of this protease, R-X-(L/I/V)-X↓ is different from that of furin R-X-(K/R)-R↓ [23]. However, inspection of the furin motifs in filovirus GPs revealed an overlapping SKI-1 motif in the GPs of Marburg-, Ebola- and Cuevaviruses (Fig 1), which likely supports MARV-GP cleavage by SKI-1. This motif is not intact in EBOV-GP Mayinga and both EBOV-GP Mayinga and Makona harbor an glutamate residue at P3 and an isoleucine residue at P1 (cleavage occurs between residues P1 and P1’), which are not found in several other viral and cellular SKI-1 substrates [64]. One can thus speculate that these residues might account, at least in part, for the absence of EBOV-GP Makona cleavage by SKI-1 in the present study.

We addressed whether determinants other than compromised proprotein convertase activity might contribute to the reduced MARV-GP cleavage in *SEC61B*-KO cells. This line of investigation was triggered by our finding that a faint smear rather than a discrete GP2 band is detectable for MARV-GP in *SEC61B*-KO cells and that this smear is due to the presence of heterogenous GP2 glycoforms. Published studies indicated that oligomannose N-glycans are attached to EBOV-GP2 residue N563 [65] and that mutation of this residue markedly reduces GP cleavage [53]. Similarly, mutation of N618, which is decorated with paucimannose N-glycans [65], slightly reduced EBOV-GP cleavage [53], suggesting that appropriate N-glycosylation of EBOV-GP2 is important for EBOV-GP cleavage. Therefore, we analyzed whether the same applies to MARV-GP. Mutation of N564 (the equivalent of N563 in EBOV-GP) jointly with PNGaseF digest revealed that N564 is N-glycosylated and that the integrity of this residue is essential for robust MARV-GP cleavage and MARV-GP-driven entry (Fig 7). In contrast, residue N619, although being decorated with N-glycans, was largely dispensable for MARV-GP cleavage and GP-mediated cell entry. Further, we confirmed that residue N563 is required for EBOV-GP cleavage, although this effect was detected mainly with VSV particles and not cell lysates, and showed that mutation of N563 has only a minor effect on EBOV-GP-driven cell entry, in keeping with published data [51–53]. Collectively, these results indicate that the absence of adequate N-glycosylation of MARV-GP residue N564 in *SEC61B*-KO cells can contribute to the inefficient GP cleavage in this cell line.

Inhibition of Sec61β or other components of the Sec61 translocon might constitute a novel approach to antiviral therapy. To determine whether Sec61β is required for infectivity of filoviruses, we first employed VSV WT and VSV-chimeras encoding filovirus glycoproteins. WT VSV efficiently infected *SEC61B*-KO cells while infection with rVSV-MARV-GP and rVSV-EBOV-GP was markedly reduced, likely due to diminished particle incorporation of GP (Fig 8). This finding suggests that both MARV-GP and EBOV-GP might not be appropriately trafficked to the plasma membrane in *SEC61B*-KO cells, the site of VSV budding. In contrast, authentic EBOV Mayinga and MARV Musoke infected *SEC61B*-KO cells with only slightly reduced efficiency as compared to WT cells. This points towards differences in GP trafficking in cells infected with VSV-chimeras relative to authentic filoviruses or, more likely, to higher overall GP expression in filovirus infected cells, allowing for virion incorporation of GP at levels sufficient for robust infectivity.

Inhibitors of the Sec61 translocon can block ER import in a client specific manner [66] and are currently evaluated within phase I clinical trials for cancer treatment. Furthermore, the Sec61 translocon inhibitor Apratoxin S4 [67–69] has recently been shown to inhibit SARS-CoV-2 and influenza A virus infection [32]. Further, a preprint suggests that the Sec61 translocon inhibitor mycolactone can inhibit ER translocation of viral glycoproteins but has only limited effects on viral assembly for which expression of low amounts of viral glycoproteins is frequently sufficient [70]. Here, we show that Apratoxin S4 blocks infection with authentic EBOV and MARV in a potent and concentration dependent manner without eliciting unwanted cytotoxicity (Fig 8), indicating that this compound merits further exploration for anti-filovirus therapy.

In sum, we demonstrate that MARV-GP but not EBOV-GP cleavage is required for GP-driven cell entry, demonstrate that SKI-1 cleaves MARV-GP and provide evidence that ER import of GP might be a viable target for antiviral intervention in the context of filovirus infection.

### Limitations of the study

Results obtained with filovirus GP transfected cell lines or cell lines infected with pseudoviruses harboring GP await confirmation with primary cells and authentic filoviruses. It remains to be determined which protease(s) cleave(s) EBOV-GP at the furin motif in *SEC61B*-KO cells and why the integrity of this site is dispensable for GP-driven cell entry. Providing answers will be challenging, since a recent study reported that EBOV-GP exhibits considerable flexibility regarding choice of cleavage site and activating protease [39].

## Acknowledgements

We thank Lina Elsasser for excellent technical assistance. H.L., T.S., E.B.-F. and S.P. were supported by the European Commission, Horizon Europe, VIGILANT (101041799), H.H.-W. was supported by Deutsche Forschungsgemeinschaft (HO 5546/3-1), H.L. was supported by the Debbie and Sylvia DeSantis Chair professorship. A.-D. W. received funding from the LOEWE Center DRUID, project A1. N.G.S was supported by Canadian Institute of Health Research (CIHR) grants # 191678 and 950-231335. The funders had no role in study design, data collection and analysis, decision to publish, or preparation of the manuscript.

## Author contributions

Conceptualization: Stefan Pöhlmann and Michael Winkler

Formal Analysis: Katharina E. Decker, Anke-Dorothee Werner, Stefan Pöhlmann and Michael Winkler

Funding Acquisition: Heike Hofmann-Winkler and Stefan Pöhlmann

Investigation: Katharina E. Decker, Anke-Dorothee Werner, Heike Hofmann-Winkler, Lu Zhang, Pamela Stomberg, Sabine Gärtner, Amy Kempf, Inga Nehlmeier, and Michael Winkler

Contribution of essential reagents: Markus Hoffmann, Qi-Yin Chen, Hendrik Luesch, Torsten Steinmetzer, Eva Böttcher-Friebertshäuser, Nabil G. Seidah

Writing – Original Draft: Katharina E. Decker and Stefan Pöhlmann

Writing – Review: & Editing: All authors

## Declaration of Interests

H.L. is cofounder of Oceanyx Pharmaceuticals, Inc. that has licensed patents and patent applications related to apratoxins.

## Supporting information

**Figure S1: Synthesis of MI-2701 (related to Figure 6).**

(A) Structure of inhibitor MI-2701 (N-(2-methoxyphenethyl)-4-((3-phenylpiperidin-1-yl)methyl)- N-(pyrrolidin-3-yl)benzamide x 2 TFA).

(B) Inhibitor **MI-2701** was synthesized as shown in Scheme S1 from intermediate **1**, which was prepared as described previously (Hay et al., 2007). Briefly, the aldehyde derivative **1** (50 mg, 0.11 mmol, 1.0 equiv) was dissolved in 2 mL dry methanol in presence of molecular sieve 4A and treated with 3-phenylpiperidine **2** (24 mg, 0.15 mmol, 1.4 equiv). The mixture was stirred for few hours and subsequently treated with NaBH_3_CN (10 mg, 0.15 mmol, 1.4 equiv), followed by stirring for additional 2 hours. The mixture was filtrated, the solvent evaporated in vacuo, and the remaining residue treated with 2 mL 4 N HCl in dioxane. The product was precipitated by addition of diethyl ether and isolated by centrifugation, followed by purification with preparative reversed phase HPLC. Yield: 17.8 mg (0.0312 mmol, 28.4 %) white hygroscopic solid. HPLC: 29.50 min, start at 10 % B (purity: 98.7 %). MS (ESI, positive): calcd, 497.30; *m/z* 498.48 [M+H]^+^ and 249.88 [M+2H]^2+^/2.

## Materials and methods

### Cell Culture

Human embryonic kidney (HEK293T, DSMZ, ACC 635) cells, HEK293T-derived *SEC61B*- KOcells, Vero76 (ATCC CRL-1587, kind gift by A. Maisner) cells and Huh7 (kindly provided by Thomas Pietschmann) cells were cultured in Dulbecco’s modified Eagle’s medium (DMEM, PAN Biotech) containing 10 % fetal bovine serum (FBS, PAN Biotech) 100 U/mL Penicillin and 0.1 mg/mL Streptomycin (P/S, Pan Biotech, P06-07100) in a humidified atmosphere at 37 °C and 5 % CO_2_. For splitting and seeding, cell lines were washed with phosphate-buffered saline (PBS, in-house production) and detached with trypsin/EDTA solution (0.05 %/0.02 % in DPBS w/o Ca/Mg; PAN Biotech, P10-023100). Cells were routinely tested for mycoplasma contamination.

### Generation of CRISPR/Cas9 knockout cell lines

For generation of *SEC61B*-KOcells, a *SEC61B* targeting sgRNA (5’- GGGCGGCGGGATCCACTGTC-3‘) selected from the GeCKOv2 library was cloned into the pLentiCRISPRv2 vector using Esp3I restriction sites [71, 72]. The plasmid containing the *SEC61B*-targeting sgRNA was then transfected into HEK93T cells using the calcium-phosphate precipitation method. The next day the cell culture medium was replaced with fresh DMEM containing 10 % FCS, P/S and supplemented with puromycin (1 µg/mL, PAA) to select for transfected cells. After 3 days, cells were seeded with a density of 1 cell/well in DMEM/F12 (+10% FBS, +P/S) into 96-well plates to select single cell clones. To promote growth, filtered, HEK293T-conditioned medium was added (diluted 1:2 with fresh medium). Absence of Sec61β expression in single cell clones was confirmed via immunoblot.

### Plasmids

Vector pLentiCRISPR v2 was a gift from Feng Zhang (Addgene plasmid #52961; [71]). To generate a Sec61β expression plasmid, a codon-optimized and sgRNA resistant *SEC61B* Gene String sequence was designed (containing a 5’ XhoI and a 3’ EcoRI restriction site) and ordered as Gene String from Thermo Fisher Scientific. This fragment was amplified via PCR and was subsequently cloned into a pCAGGS vector via EcoRI and XhoI. pCAGGS empty vector or encoding for the GPs (with and without V5 tag) of Indiana vesiculovirus (VSV, ABD73123.1), Ebola virus (EBOV strain Makona, AIE11800.1; EBOV strain Mayinga, NP_066246.1; V5 tagged plasmids of Makona and Mayinga strain carry the I662V mutation), Sudan virus (SUDV, ACR33190.1), Bundibugyo virus (BDBV, YP_003815435.1), Taï Forest virus (TAFV, YP_003815426.1), Reston virus (RESTV, Q66799.1), Marburg virus (MARV, strain Musoke, YP_001531156.1); MARV bat2007 (Uganda 371Bat2007, ACT79236.1), Raven virus (RAVV; Uganda 982Bat2008, ACT79222.1), Lloviu virus (LLOV, UJP71058.1, kind gift from A. Takada, Hokkaido University Research Center for Zoonosis Control, Sapporo, Japan), Lymphocytic choriomeningitis virus (LCMV, strain clone 13, ABC96001.2) and Lassa virus (LASV, strain Josiah, NP_694870.1) were described elsewhere [73–76]. To generate mutants having a defective furin cleavage site (dFCS) for GPs of all *Orthoebolavirus*, *Orthomarburgvirus* and *Cuevavirus* species, for all GPs primers were designed to mutate all basic amino acids (arginine and lysine) of the furin cleavage site to alanine. Additionally, for MARV-GP a mutants already described by Volchkov and colleagues (RRKL^435^, [25]) and a mutant containing a monobasic cleavage site (AAAR^435^) were generated. Mutated GPs were generated and amplified by hybrid PCR and cloned into the pCAGGS vector via restriction enzymes. pCAGGS-EBOV-GP(Makona) and pCAGGS-EBOV-GP-V5(Makona) with either N563D, N618D or both mutations as well as pCAGGS-MARV-GP-V5(Musoke) with either N564D, N619D or both mutations were cloned by using specific mutagenesis primers, amplification of the GP by hybrid PCR and cloning into the pCAGGS vector via EcoRI and NheI. For generation of pQCXIP_hFurin-cMYC, RNA from A549 cells (RNeasy Kit, Qiagen) was isolated, cDNA was synthesized (SuperScript III First-Strand Kit, Thermo Fisher Scientific) and hFurin was amplified via PCR, introducing a NotI and an EcoRI restriction site. Upon digestion with NotI and EcoRI, hFurin was ligated into a pQCXIP vector (Clontech, Palo Alto) modified to contain a multiple cloning site. pIR-eGFP, pIR-hPCSK7-eGFP-V5, pIR-hPCSK9-eGFP-V5, pIR-PACE4-eGFP-V5, pIR-hSKI-1-eGFP-V5, pIR-mPC5A-eGFP-V5 were described elsewhere [35, 77–81]. The V5-tag was exchanged by a cMyc-tag by introducing restriction sites after the coding sequence of the proprotein convertases with overhang primer (NotI for PCSK7, PCSK9 and PACE4, BamHI for SKI-1 and MluI for mPC5A). After PCR amplification, cMyc-tagged PCs were inserted into the same vector (pIRES2-EGFP) via restriction digest. All plasmids were checked for integrity via Sanger sequencing (MicroSynth, Göttingen). Detailed cloning strategies, primer and plasmid sequences can be shared upon request.

### CFSE Assay

To compare cell growth between HEK293T and HEK293T-derived *SEC61B*-KOcells, CFSE staining (CFSE Cell Division Tracker Kit, BioLegend, 423801) was performed. For this, 1×10^6^ cells were incubated in 1 mL PBS with 5 µM CFSE or PBS only as control for 20 min at 37 °C in the dark. The staining reaction was stopped by adding 6 mL of DMEM with 10 % FCS and P/S and centrifuged. After resuspension in fresh DMEM (+ 10 % FCS and P/S) cells were seeded in a 12-well plate with 1 mL/well and a density of 1x 10^5^ cells/mL and incubated at 37°C with 5% CO2. After 0, 1, 2 and 3 days, cells from CFSE and control treated samples were harvested for flow cytometric analysis.

### Transfection

For transfection, cells were seeded in DMEM/F-12, GlutaMAX^TM^ medium (Fisher Scientific, Gibco^TM^, 31331093) supplement with 5% FBS. HEK293T cells were seeded with a cell density of 1.2 x 10^5^ cells/mL per well whereas *SEC61B*-KOcells were seeded with a density of 1.4 x 10^5^ cells/mL in 12-well or 6-well cell culture plates with 1 mL/well or 2 mL/well, respectively. One day later, HEK293T and *SEC61B*-KOcells were transfected by using the polyethylenimine (PEI) transfection method as described by Longo and colleagues [82]. In brief, for one well of a 12-well cell culture plate 1 µg plasmid DNA and 3 µL of PEI (Polyscience, Inc., 23966, 0.1 % w/v in water) solution were diluted in 50 µL Opti-MEM (Fischer Scientific, Gibco^TM^, 15392402) each. Then, diluted DNA and PEI were mixed, incubated for 30 min at RT and added carefully to the side of the well. For cotransfection of GP with PC plasmids, 0.7 µL of GP-expressing plasmids and 0.4 µL of PC-expressing plasmids were used. For cotransfection of Sec61β with GP plasmids, 0.5 µg per plasmid and well were transfected.

### Immunoblot

For analysis of protein expression in whole cell lysates (WCL) by immunoblot, cells were seeded in 12-well cell culture plates and transfected one day later as described above. Three days after transfection cell culture supernatant was removed and cells were harvested by adding 100 µL of 1x SDS-containing lysis buffer (62.5 mM Tris [pH 6.8], 10 % glycerol, 2 % SDS, 1 mM EDTA, 0.005% bromphenol blue, 5 % β-mercaptoethanol) per well. After 10 min incubation at room temperature, samples were boiled at 95 °C for 10 min and stored at −20 °C until further use. Samples were separated on SDS-polyacrylamid gels (12.5% acrylamid) and blotted onto nitrocellulose membranes (Cytiva, 10600001) if not otherwise specified or PVDF (Cytiva, 10600023). PVDF membranes were activated before usage for 45-60 s in 100% methanol, then washed in distilled water. After blotting, membranes were blocked for 1 h at room temperature (RT) in 5% skim milk in PBS containing 0.1 % Tween-20 (PBS-T). Membranes were then incubated with primary antibodies for either 1 h at RT or overnight at 4 °C. Primary antibodies were diluted in 5% skim milk in PBS-T, except the β-actin specific antibody (Sigma-Aldrich, A5441, mouse, 1:500) which was used for the loading control of WCL which was diluted in 5% BSA in PBS-T. Besides β-actin, GAPDH visualization (Proteintech, 60004, mouse, 1:5000) served as loading control. Equal loading of VSVpp was assessed by a VSV-M specific antibody (Kerafast, EB0011, mouse, 1:1000). V5-tagged GPs from WCL and VSVpp were visualized using a V5-specific antibody (Invitrogen, mouse, R960-25, 1:5000). The proprotein convertases (PCs) were stained either with antibody targeting the cMyc epitope (antibodies-online, ABIN559687, mouse, 1:1000) or with specific antibodies against furin (Invitrogen, PA1-062, rabbit, 1:1000), and SKI-1 (polyclonal: GeneTex, GTX117475, rabbit, 1:500 or monoclonal: Santa Cruz Biotechnology, sc-271916, mouse, 1:1000). Sec61β was detected with a Sec61β-specific antibody (Proteintech, 51020-2-AP). After incubation with the primary antibody, the membranes were washed three times with PBS-T and then incubated either with mouse specific (Dianova, 115-035-003, 1:1000) or rabbit specific (Dianova, 111-035-004, 1:1000) horseradish peroxidase (HRP)-conjugated antibody for 1 h at RT. Finally, membranes were washed three times with PBS-T and signals were developed using enhanced chemiluminescence (ECL) substrate (Cyanagen, XLS1420250) and recorded with the ChemoCam imaging system (Intas) using the ChemoStarProfessional software (Intas). For consecutive staining, membranes were washed after imaging with PBS-T and stripped with stripping buffer (Carl Roth, Roti-Free stripping buffer 2.0, 3319.2 or 2.2 plus, 3337.2) by shaking for 1h at RT. After washing thoroughly four times with PBS-T membranes were blocked, stained and developed as described before.

### PNGase F digest of cell lysates

GP transfected HEK293T or *SEC61B*-KOcells were harvested and washed with 1 mL PBS and then divided into two tubes. One sample was digested with PNGase F (New England Biolabs, P0704) according to the manufacturer’s instructions for denaturing reaction conditions (but with 2 instead of 1 µL of PNGase F), while the control sample was treated with water instead of PNGase F. After digestion, 22 µL 2x SDS-containing lysis buffer were added and samples were prepared for immunoblot.

### Production of rhabdoviral pseudo particles

To analyze GP driven entry into target cells, single cycle rhabdoviral reporter particles were generated as described [83, 84]. HEK293T and *SEC61B*-KOcells were transfected with GP expressing plasmids as described above. Two days after transfection the cell culture medium was removed and cells were infected with VSV*ΔG(eGFP,fLuc) (kindly provided by Gert Zimmer, [84]) trans-complemented with VSV-G at MOI 3. For this, VSV*ΔG(eGFP,fLuc)_VSV-G was diluted in DMEM (10 % FCS, P/S) and added to the cells in reduced volume (0.5 mL for 12-well and 1 mL for 6-well plates). After 1 h at 37 °C the inoculum was removed and fresh DMEM (10 % FCS, P/S) containing VSV-G specific antibodies (1:1000, hybridoma cell supernatant, ATCC CRL-2700) was added to the cells (1 mL for 12-well and 2 mL for 6-well plates) to neutralize remaining VSV*ΔG(eGFP,fLuc)_VSV-G. The cell culture supernatant was harvested 18-20 h after infection. Cellular debris was removed by centrifugation at 2000 x g for 10 min and clarified supernatant was used either directly or stored at −80 °C until use.

### Analysis of GP incorporation into VSV particles

To analyze GP incorporation into VSV particles, 1.3 mL pseudotyped particles cleared from cellular debris were transferred into a 1.5 mL reaction tube. 50 µL of sucrose (20 % w/v in PBS) was added carefully to the bottom of the reaction tube to create a sucrose cushion. After centrifugation at 16.000 x g for 90 min at 4°C, the cell culture supernatant was removed while leaving the colorless, VSV particle containing sucrose cushion undisturbed. After this, the sucrose cushion was mixed with 50 µL of 2x SDS-containing lysis buffer. The samples were boiled at 95 °C for 10 min and either stored at −20 °C or directly loaded onto a SDS-PAGE.

### Infection experiment with rhabdoviral pseudotypes

HEK293T, Huh7 and Vero76 cells were seeded in 96-well cell culture plates at a cell density of 2×10^5^ cells/mL (for HEK293T and Huh7 cells) or 1x 10^5^ cells/mL (for Vero76 cells) in 100 µL/well DMEM (10 % FCS and SP) and incubated at 37 °C. The next day, the cell culture medium was removed and replaced with 100 µL/well supernatant containing pseudoparticles. After incubation for 18-20 h at 37 °C, the culture supernatants were removed and cells were lysed in 50 µL PBS containing 0.1 % triton-X-100 for 20-30 min at RT. Cell lysates were then transferred into opaque 96-well plates. Finally, 50 µL/well firefly luciferase substrate (Beetle Juice, PJK, 102511) was added to the cell lysates and luciferase activity was measured using a Hidex Sense Microplate Reader.

### Production and infection experiments with VSV chimeras

For production of replication-competent, chimeric VSV expressing EBOV-GP, MARV-GP, or LASV-GPC instead of VSV-G, a plasmid-encoded VSV anti-genome (GenBank: J02428.1) was modified as follows. First, unique MluI and NheI restriction sites were introduced upstream and downstream of the VSV-G gene to facilitate subsequent modifications. Next, a genetic cassette flanked by MluI and NheI restriction sites and consisting of VSV-G and eGFP open reading frames, which were separated by a minimal intergenic region [85] with unique AscI and NotI restriction sites (<u>GGCGCGC</u>CCTCAAATCCTGCTAGGTATGAAAAAAACTAACAGATATCAC<u>GCGGCCGC</u>, underlined letters indicate restriction sites), was generated by overlap-extension PCR and used to replace the original VSV-G open reading frame. This resulted in plasmid pVSV* (the asterisk indicates the presence of an additional transcription unit for eGFP in the VSV genome). Finally, the VSV-G open reading frame of pVSV* was replaced by open reading frames for EBOV-GP (strain Mayinga), MARV-GP (strain Musoke), or LASV-GPC (strain Josiah), making use of MluI and AscI restriction sites. This procedure resulted in plasmids pVSV*ΔG (EBOV-GP), pVSV*ΔG (MARV-GP), and pVSV*ΔG (LASV-GPC). The integrity of all PCR-amplified sequences was verified by automated sequence analysis using a commercial service (MicroSynth, Göttingen). Rescue of replication-competent, chimeric VSV was performed as described before [86].

Replication kinetics of recombinant, and eGFP encoding chimeric viruses in HEK293T and *SEC61B*-KOcells (clone 7) was determined. For this, HEK293T and *SEC61B*-KO cells were seeded in 48-well cell culture plates. After one day, cells were infected with rVSV, rVSV-MARV-GP and rVSV-EBOV-GP with 0.01 MOI to allow multiple cycles of infection. After infection for 1 h, inoculum was removed and replaced with fresh DMEM (2 % FCS, P/S). At specific time points, cells were harvested and prepared for flow cytometry analysis (0, 8, 24, 32, 48, 56, 72 hours post infection, hpi) to quantify eGFP-expressing infected cells, or supernatant was harvested (0, 24 and 48 hpi) and stored at −80 °C until viral load was determined by TCID_50_.

Median tissue culture infectious dose (TCID50) was determined to measure the titer of rVSV chimera stocks (titrated on Vero76 or HEK293T, as indicated) and to determine the viral titer in the supernatant of infected cells (titrated on Vero76). For this, the cell culture supernatant was harvested at indicated time points, centrifuged at 2000 x g for 5 min to remove cells and stored at −80 °C until titration was performed. For titration, the supernatant was diluted in DMEM supplemented with 2.5 % FBS and P/S in a 10-fold serial dilution (from 10^-1^-10^-12^) and then added in quadruplicates onto Vero76 or HEK293T cells. Infected wells were identified by detection of eGFP fluorescence (recombinant VSV chimera) or cytopathic effect (authentic EBOV and MARV). Calculation of viral titers was conducted according to the Spearman and Kärber method (TCID_50_ calculation spreadsheet from Marco Binder; calculation as described in [87]).

### Flow cytometry

Flow cytometry was used to measure CFSE signals and to determine the percentage of cells infected with eGFP encoding chimeric VSV. For this, cells were harvested at the indicated time points by resuspending them in cell culture medium followed by washing with PBS containing 5 mM EDTA and a second washing step with FACS buffer (PBS with 2 % BSA, 2 mM EDTA, 0.2 % sodium azide). Afterwards, cells were resuspended and fixed with a 3:1 mixture of 4 % PFA solution and FACS buffer. Samples were stored at 4 °C until measurement, but not longer than three days. Measurement was performed using the ID7000 Spectral Cell Analyzer ID7000 (Sony). Data were analyzed either with ID7000 Software (VSV chimera infections, version 2.0.2.17121) or with FlowJo (CFSE assay, version 10.8.1). The living cell population was gated based on forward scatter (FSC) and sideward scatter (SSC). The gate for GFP positive cells aligned to the mock-infected control.

### Antiviral activity of protease inhibitors

HEK293T cells were seeded with 3.6×10^4^ cells/well in 48-well cell culture plates in 300 µL/well and incubated at 37 °C over night. Afterwards, cells were infected with rVSV, rVSV-LASV-GP, rVSV-MARV-GP or rVSV-EBOV-GP (MOI 0.01). After 1 h, inoculum was removed and 300 µL fresh DMEM with 10 % FCS and P/S was added. After 8 h, protease inhibitors were added. For inhibition of furin and furin-like proteases Decanoyl-RVKR-cloromethylketone (Tocris, 3501) was dissolved in water and used at a concentration of 25 µM. Inhibition of SKI-1 was investigated using PF-429242 dihydrochloride (Sigma-Aldrich, SML0667), described in [45], and MI-2701 (N-(2-methoxyphenethyl)-4-((3-phenylpiperidin-1-yl)methyl)-N-(pyrrolidin-3-yl)benzamide x 2 TFA). The chemical structure of inhibitor MI-2701 is shown in the supporting information and similar to compound 7a published by Hay and colleagues [45] but without methylation on the 3-(phenyl)piperidine segment. Both inhibitors were dissolved in DMSO and used at a concentration of 5 µM (PF-429242) or 1 µM (MI-2701). After 0, 8, 24, 32, 48, 56, and 72 h post infection cells were harvested and prepared for flow cytometry analysis. For investigation of protease inhibitors on GP cleavage efficiency, HEK293T cells were transfected with MARV-GP plasmid. After 8 h, the medium was removed and different concentrations of the furin inhibitor decanoyl-RVKR-CMK (0.1, 1, 10, and 25 µM; H_2_O as control) and the SKI-1 inhibitor PF-429242 (0.01, 0.1, 1, and 5 µM; DMSO as control) were added. Following an incubation of two days at 37 °C, cells were harvested and prepared for immunoblot analysis.

### Antiviral activity of Apratoxin S4

Apratoxin S4 was produced as previously described [68]. For analysis of Apratoxin S4 antiviral activity, HEK293T and Vero76 cells were seeded in 96-well cell culture plates with 100 µL/well and a density of 3×10^5^ or 1×10^5^ cells/well, respectively and incubated at 37 °C. The next day, cell culture medium was removed and replaced with DMEM (+ 10 % FCS and P/S) containing 1, 0.1 or 0.01 µM Apratoxin S4 or DMSO as control and incubated for 2 h at 37 °C. Then, recombinant chimeric viruses were added to the Apratoxin S4 treated cells with a MOI of 0.01. After 24 h (for rVSV) or 48 h (for rVSV-MARV-GP and rVSV-EBOV-GP) cells were collected and prepared for flow cytometry analysis.

### Cell Viability Assay

Cell viability was assessed by determining cellular ATP level with the CellTiter-Glo® Luminescent Cell Viability Assay (Promega, G7571) by following the manufacturer’s instructions.

### BSL-4 Experiments

All infection experiments using Ebola virus (EBOV, strain Mayinga) and Marburg virus (MARV, strain Musoke) were performed in the BSL4 laboratory of the Institute for Virology, University of Marburg, according to national and international regulations.

HEK293T and *SEC61B*-KOcells were seeded in 6 well cell culture plates (4×10^5^ and 8×10^5^ cells, respectively, approx. 50 % confluency) and infected with EBOV and MARV. For this, the cell culture medium was aspirated and replaced by 1.5 mL DMEM (3 % FCS) containing MARV or EBOV with a MOI of approx. 0.1. 1, 24 and 48 hpi, 250 µL of the supernatant were harvested.

Vero76 cells were seeded in 24-well plates to a confluency of 50 % and treated with Apratoxin S4 in varying concentrations (1-10^-2^ µM) or DMSO (0.1 % v/v) in 200 µL DMEM (3 % FCS). After incubation for 2 h at 37 °C, cells were infected with EBOV or MARV at an MOI of 0.1 by adding the required virus volume directly to the supernatant. The supernatant was collected either directly after the infection (input control, 0 hpi) or after 2 dpi. Viral titers of the harvested samples were assessed by titration (TCID_50_ method) and calculated using the Spearman and Kärber algorithm.

### HPLC and ESI spectrometry

For production of MI-2701, analytical reversed-phase HPLC measurements were performed on a Primaide system (VWR, Hitachi, column: NUCLEODUR C18 ec, 5 μm, 100 Å, 4.6 mm x 250 mm, Macherey-Nagel, Düren, Germany) with 0.1 % TFA in water (solvent A) and 0.1 % TFA in acetonitrile (solvent B) as eluents using a linear gradient with an increase of 1 % solvent B/min at a flow rate of 1 mL/min and detection at 220 nm. Purifications via preparative reversed-phase HPLC were performed on a Knauer Azura system (pump P 2.1 L equipped with pump head E4099AB, detector UVD 2.1L, Knauer GmbH, Berlin, Germany, column: NUCLEODUR C18 ec, 5 μm, 100 Å, 32 mm x 250 mm, Macherey-Nagel, Düren, Germany) using the same solvents as described above for analytical HPLC and a linear gradient with an increase of 0.5 % solvent B/min at a flow rate of 20 mL/min (detection at 220 nm, in few cases with larger amounts at 254 nm). After preparative HPLC, the inhibitors were obtained as lyophilized TFA-salts in a purity > 95 % using a freeze-drier (Martin Christ GmbH, Osterode am Harz, Germany).

ESI mass spectra were measured on a QTrap 2000 ESI spectrometer (Applied Biosystems, now part of Thermo Fischer Scientific).

### Light Microscopy

Bright field images were taken on a LSM800 (Zeiss) with a 20x objective and an electronically switchable illumination and detection module (ESID). Images were recorded with the ZEN 2.3 imaging software (Zeiss) and prepared for publication with the Zeiss Zen 3.8 lite software (Zeiss).

### Statistical Analysis

Statistical analysis was performed using GraphPad Prism 10 software (version 10.4.1). Statistical tests used for the respective data sets are indicated in the figure legends.

