## Supplementary Figure S1 for "Proteolytic processing of the Marburg virus glycoprotein depends on Sec61β and is required for cell entry"

### Supplemental Figure S1

### A)

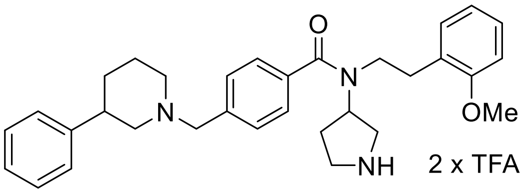


### B)

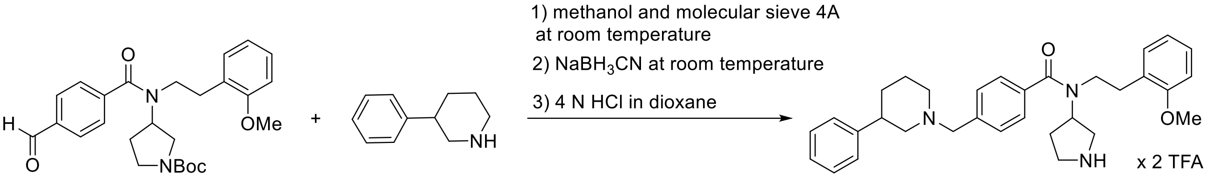


**Supplemental Figure S1. Synthesis of MI-2701**

(A) Structure of inhibitor MI-2701 (N-(2-methoxyphenethyl)-4-((3-phenylpiperidin-1-yl)methyl)-N-(pyrrolidin-3-yl)benzamide x 2 TFA).

(B) Inhibitor **MI-2701** was synthesized as shown in Scheme S1 from intermediate **1**, which was prepared as described previously (Hay et al., 2007). Briefly, the aldehyde derivative **1** (50 mg, 0.11 mmol, 1.0 equiv) was dissolved in 2 mL dry methanol in presence of molecular sieve 4A and treated with 3-phenylpiperidine **2** (24 mg, 0.15 mmol, 1.4 equiv). The mixture was stirred for few hours and subsequently treated with $NaBH_3CN$ (10 mg, 0.15 mmol, 1.4 equiv), followed by stirring for additional 2 hours. The mixture was filtrated, the solvent evaporated in vacuo, and the remaining residue treated with 2 mL 4 N HCl in dioxane. The product was precipitated by addition of diethyl ether and isolated by centrifugation, followed by purification with preparative reversed phase HPLC. Yield: 17.8 mg (0.0312 mmol, 28.4 %) white hygroscopic solid. HPLC: 29.50 min, start at 10 % B (purity: 98.7 %). MS (ESI, positive): calcd, 497.30; *m/z* 498.48 $[M+H]^+$ and 249.88 $[M+2H]^{2+}/2$.
